# Fat body driver expression report across *Drosophila melanogaster* tissues and sex

**DOI:** 10.64898/2026.05.29.728847

**Authors:** IH Williams, ME Meyerink, Alissa R Armstrong

## Abstract

*Drosophila melanogaster* serves as a valuable model system for advancing our understanding of adipose tissue given its analogous organ systems, conserved metabolic, endocrine, and nutrient-sensing functions, and well-established genetic tools. Among the most widely used genetic tools is the *Gal4/UAS* system. Several *Gal4* driver lines are reported to control expression in the *D. melanogaster* fat body, but secondary expression sites and responses to physiological changes have not been fully characterized. In this report, we describe fluorescent reporter expression for 31 fat body *Gal4* transgenic lines in larvae and adults, males and females, and in multiple tissues, brain, indirect flight muscle, gut, ovary, testis, and fat body. This screen highlights expression differences with respect to level, sex, and stage. Moreover, we find that several lines drive expression in tissues that had not been previously described. Together, we present a comprehensive expression atlas of fat body driver lines, serving as a resource to advance adipose tissue research.

## Introduction

Historically viewed as inert, adipose tissue functions as a central metabolic and endocrine organ (Harvey et al., 2020). In recent decades, the growing obesity epidemic, along with associated comorbidities, such as type 2 diabetes, cardiovascular and liver diseases, and certain cancers, has driven research on adipose tissue (Corvera et al., 2026). While rodent models are routinely used to investigate adipose tissue dysfunction associated with metabolic disorders (Lutz & Woods, 2012; Martins et al., 2022), the versatile model organism *Drosophila melanogaster* offers an efficient, lower cost means to further our understanding of adipose tissue function in physiological and pathophysiological states. Analogous to white mammalian adipose tissue and liver, the adult *D. melanogaster* fat body is primarily localized to the abdomen, where it consists of nutrient-storing adipocytes and hepatocyte-like oenocytes (Huang & Perrimon, 2026). The *D. melanogaster* fat body has been studied for its involvement in energy storage, inter-organ communication, and immune response (Zheng et al., 2016; Skowronek et al., 2021). For example, the larval fat body is essential for systemic growth during development and for accumulating nutrients that are mobilized during the nonfeeding pupal stage (Arrese & Soulages, 2010). Additionally, the fat body acts as a nutrient sensor, communicating nutrient status to other tissues. Most notably, dietary input regulates activity of highly conserved nutrient-sensing pathways, like insulin/insulin-like growth factor and mechanistic Target of rapamycin signaling, in the fat body to relay nutrient status to insulin-producing cells in the brain (Géminard et al., 2009) and the ovarian germline stem cell lineage (Armstrong et al., 2014; Armstrong & Drummond-Barbosa, 2018; Bradshaw et al., 2024).

*D. melanogaster* as a model organism offers both practical and genetic benefits. Practical benefits include their fast life cycle, approximately 10 days from egg laid to adulthood at 25°C (Hales et al., 2015), and robust reproduction, with a single female laying hundreds of eggs over their lifespan (Drummond-Barbosa & Spradling, 2001). Genetic benefits include several well-established binary expression systems, such as Gal4/UAS, LexA/LexAop, and Q systems. These binary expression systems are available with many ready-made transgenic lines that allow tissue- and cell-type-specific, as well as temporal, control of gene expression (Zirin et al., 2024). With the capacity to perform RNA interference, overexpression, mosaic analysis with a repressible cell marker (MARCM), and genome editing using CRISPR/Cas9, all in a tissue-specific manner (Zirin et al., 2024; Lee & Luo, 1999; Port et al., 2014), binary expression systems enable researchers to genetically manipulate the fat body to assess gene, protein, and cellular functions.

One of the most widely used binary expression systems is the *Gal4/UAS* system, originally described by Brand and Perrimon in 1993 (Brand & Perrimon, 1993). In this system, the *Gal4* transgenic line, or driver, encodes the yeast transcription factor *Gal4* downstream of a promoter or enhancer region. The *UAS* transgenic line, or responder, contains several tandem repeats of an Upstream Activating Sequence (*UAS*) followed by a transgene of interest (Brand & Perrimon, 1993). Using standard genetic crosses of *Gal4* and *UAS* lines, the resultant progeny will express the transgene in select tissues or cell types dictated by the promoter or enhancer region present in the driver line. Many Gal4 lines have been reported to drive expression in the fat body, but the range of expression levels, as well as differences due to life stage and sex, have not been thoroughly described. Moreover, the expression specificity of some *fat body-Gal4* transgenic lines has not been fully reported. The goal of this report is to create a comprehensive atlas of *Gal4* lines that drive expression in the *D. melanogaster* fat body.

Using FlyBase and previous reports, we identified 31 *Gal4* lines reported to drive expression in the larval and/or adult fat body. We used a fluorescent reporter responder line to characterize expression over four criteria: 1) larval versus adult, 2) female versus male, 3) across several tissues, and 4) intensity. Overall, we found expression differences across tissues among *Gal4* lines and identified lines with stage- and sex-specific expression patterns.

## Materials and Methods

### *Drosophila* strains and culture conditions

*Drosophila* stocks were maintained at 22-25°C on standard medium containing cornmeal, molasses, yeast, and agar (Archon Scientific). Fat body *Gal4* lines screened in this study are listed in Table 1. The previously described *UAS*-controlled fluorescent reporter, *UAS-2xEGFP* (Bloomington Drosophila Stock Center, #6874), was crossed to each *Gal4* line (Halfon et al., 2002). Standard genetic crosses of *Fat body-Gal4* and *UAS-2xEGFP* lines were set to obtain progeny containing both transgenes, *Fat body-Gal4 > UAS-2xEGFP*. To assess GFP expression within larval fat body, wandering third instar larvae were collected from the side of vials. To assess GFP expression within adult tissues, adult progeny were fed the standard medium supplemented with wet yeast paste for seven days prior to dissection.

**Table 1.**
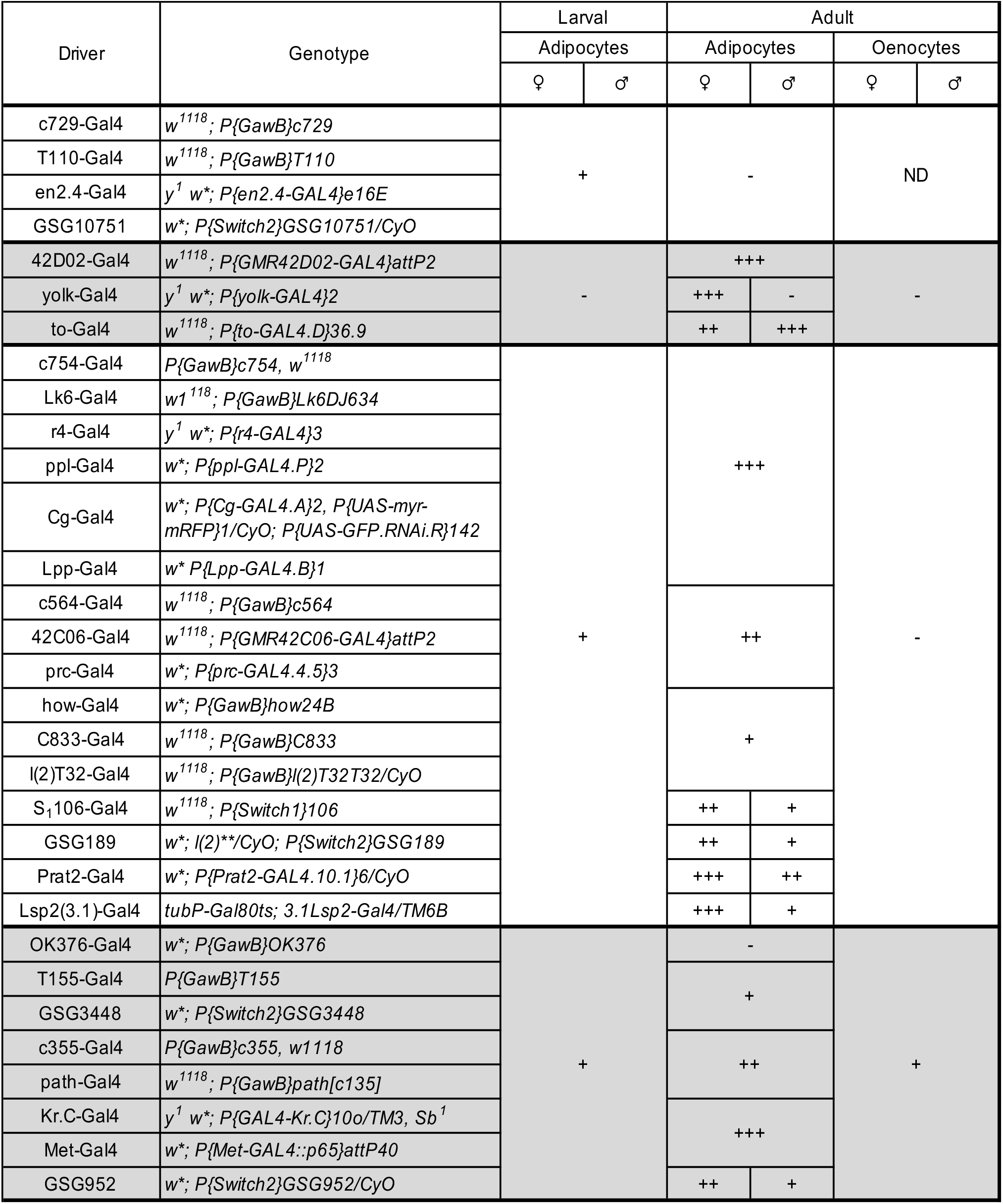
Driver fat body expression in larvae and adults.

### Temperature- and ligand-dependent induction of *Gal4*

To screen expression patterns of the temperature-sensitive driver line, tubP-Gal80ts; 3.1Lsp2-Gal4/TM6B (Lsp2(3.1)-Gal4), larval and adult progeny (Lsp2(3.1)-Gal4 > UAS-2xEGFP) were maintained at 29°C for the duration of the experiment. Larvae and adults containing the GeneSwitch Gal4 construct driving UAS-2xEGFP were fed mifepristone (RU486) (Thermo Fisher Scientific) to induce GFP expression. To feed larvae RU486, 250 µL of 1 mM RU486 in ethanol was added directly to the surface of the fly food medium 4 days post-egg-laying. RU486 was delivered to adult flies by feeding them with wet yeast paste supplemented with 1mM RU486 (1:3 vol/vol).

### Whole-mount immunostaining of tissues

All tissues were dissected in 1X phosphate saline buffer (PBS) and fixed in 5.3% paraformaldehyde diluted in 1X PBS at room temperature. Abdominal carcasses containing the fat body were dissected and pinned using the methods previously described in Ott et al., 2024. Abdominal carcasses, male reproductive tissues, and brains were fixed for 20 minutes, ovaries were fixed for 13 minutes, thoraces were fixed for 30 minutes, and guts were fixed for 1 hour. After fixation, samples were rinsed twice and washed three times for 10 minutes in either 0.1% Triton X-100 or Tween 20 diluted in 1X PBS. After fixation and washing, the female and male reproductive tissue samples underwent the following immunostaining steps, while all other tissue samples were placed in Vectashield (Vector Labs) containing DAPI immediately afterward. To add a counter label, the female and male reproductive tissue samples were incubated in blocking solution (5% bovine serum albumin, 5% normal goat serum, and 0.1% Triton X-100) overnight. Reproductive tissue samples were then incubated overnight in primary antibody solution consisting of 0.5 µg/mL mouse anti-Lamin C (LamC; Developmental Studies Hybridoma Bank (DSHB)) and 1 µg/mL mouse anti-alpha spectrin (3A9; DSHB). Following incubation in primary antibody solution, samples were washed three times for 10 minutes, incubated for 2 h with goat anti-mouse Alexa Fluor 568 (Thermo Fisher Scientific; 1:250) protected from light, and washed three times for 15 minutes. Reproductive tissue samples were then placed in Vectashield containing DAPI.

### Confocal microscopy and image acquisition

Images were acquired using a Zeiss LSM 800 confocal microscope running ZEN 2.6 software. Microscope acquisition settings were adjusted for each image to optimize visualization of GFP expression. Images of the larval fat body, adult fat body, and muscle were acquired at 20x magnification. Images of the ovariole and male reproductive component were acquired at 10x magnification, while images of the germarium were acquired at 63x magnification. Gut and brain images were acquired at 5x magnification. To qualitatively evaluate GFP levels in the fat body, GFP expression was scored manually from multiple observations of fat body patches through the eyepieces. Tissues positive for GFP expression were identified by examining multiple tissues from different flies under the eyepieces and using the Zen software.

## Results

### Larval- and adult-specific driver expression in the fat body

To characterize Gal4 line expression in the fat body and across tissues, *Gal4* lines were crossed to the UAS-controlled fluorescent reporter *UAS-2xEGFP*. The larval and adult fat body originate from distinct cell populations (Tsuyama et al., 2023); therefore, expression observed in the larval fat body may not reflect expression in the adult fat body or vice versa. To determine if each *Gal4* driver induces expression in the larval and/or adult fat body, we assessed GFP expression at both stages in females and males. The majority of lines, 24 of 31, showed GFP expression in both the larval and adult fat body (Table 1). Four lines were larva-specific, *c729-Gal4, T110-Gal4, en2*.*4-Gal4*, and *P{Switch2}GSG10751*, and drove expression in both females and males (Fig. 1a). Three lines were adult-specific, *yolk-Gal4, to-Gal4*, and *GMR42D02-Gal4*, with yolk-Gal4 only driving expression in the female fat body (Fig. 1b). Together, our analysis shows that driver expression in larval and adult adipose tissue is not always correlated.

**Fig. 1.**
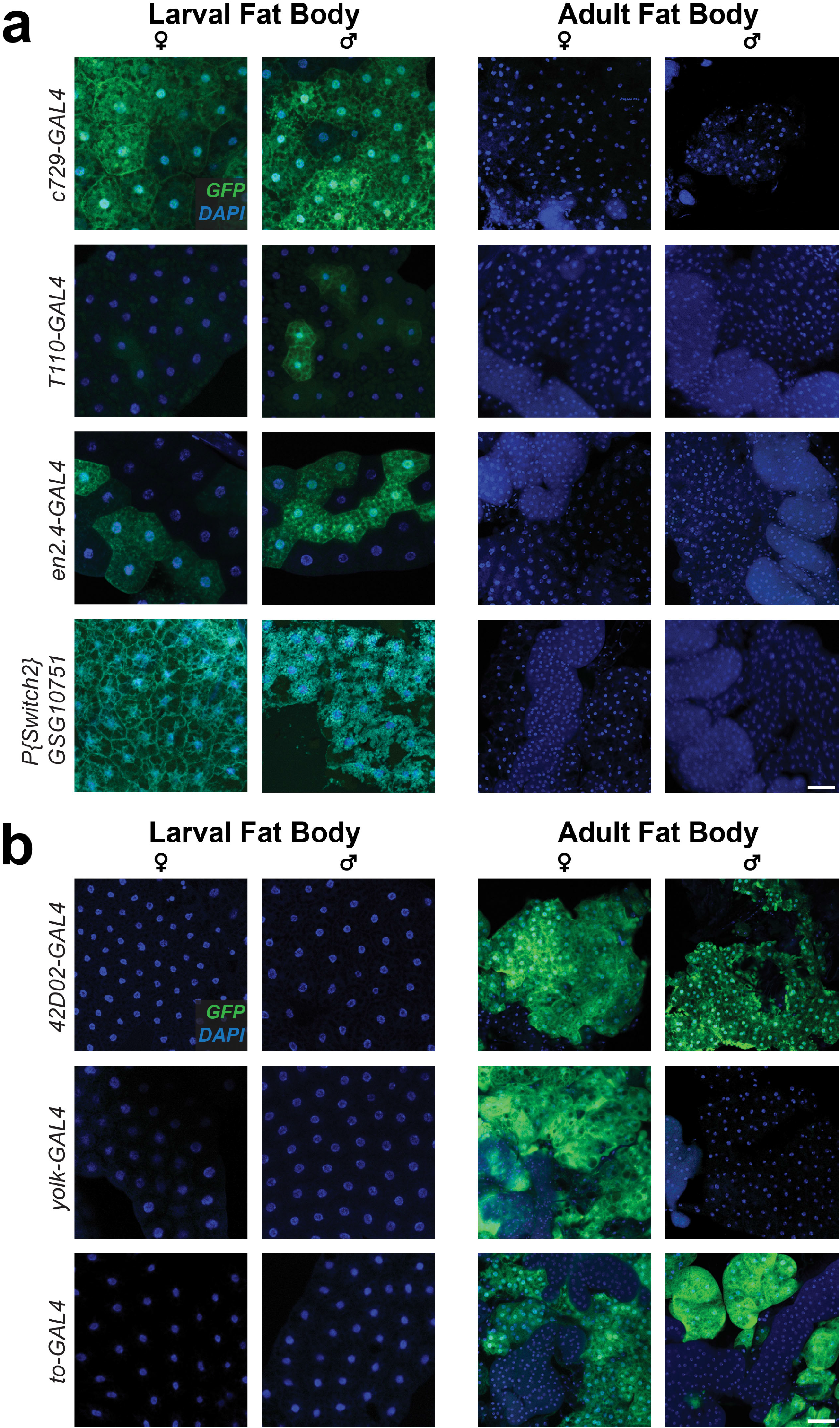
*Gal4* drivers with differences in larval and adult fat body expression. a) Lines where expression was observed in the larval fat body, but not in the adult fat body. b) Lines where expression was observed in the adult fat body, but not in the larval fat body. Endogenous GFP (green) and nuclei/DAPI (blue). Scale bars, 50 μm.

### Adult fat body drivers vary in expression level across adipocytes and oenocytes

In adults, the majority of the *D. melanogaster* fat body is located in the abdomen and consists of two cell types: adipocytes (fat cells) and oenocytes (hepatocyte-like cells) (Johnson & Butterworth, 1985). For the 27 Gal4 lines that drive expression in the adult fat body, we next asked if expression was specific to adipocytes or oenocytes or found in both cell types. We found that 17 lines induced expression in adipocytes only, one in oenocytes only, and seven in both adipocytes and oenocytes (Table 1).

Since different promoters or enhancers control each *Gal4* driver, we next asked what level of expression was induced in the adult fat body. For each line that drove expression in the adult fat body, we qualitatively scored adult adipocyte expression as weak (+), moderate (++), or strong (+++) (Fig. 2a). Out of the 26 fat body driver lines assessed, five had weak expression, five had moderate expression, and nine had strong expression in adult adipocytes for both females and males (Fig. 2b-d, Table 1). Seven lines showed sexually dimorphic expression, with *Lsp2(3*.*1)-Gal4, P{Switch}106, P{Switch2}GSG952, yolk-Gal4, P{Switch2}GSG189* and *Prat2-Gal4* showing higher expression in female than in male adipocytes, while *to-Gal4* showing slightly higher expression in male than in female adipocytes (Fig. S1-S4; Table 1). Furthermore, for *Lsp2(3*.*1)-Gal4, P{Switch}106, P{Switch2}GSG952, P{Switch2}GSG189*, and *Prat2-Gal4*, this sexual dimorphic expression is observed only in adulthood, highlighting an interaction between life stage and sex that impacts gene expression (Fig. S1-S4). Taken together, we show that different fat body *Gal4* lines induce varying levels of expression within adipocytes.

**Fig. 2.**
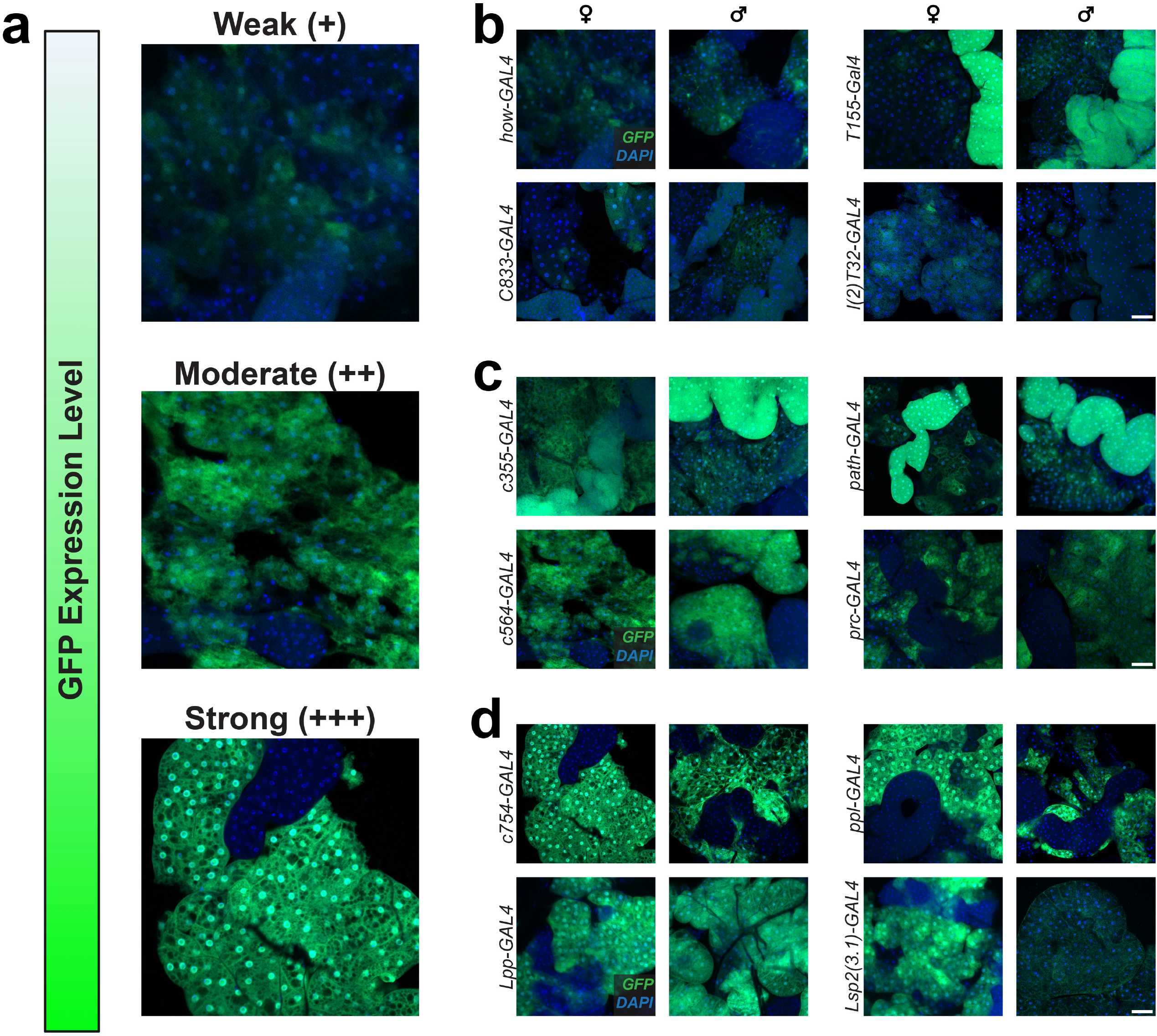
Fat body drivers ranging in expression level induced within the adipocytes. a) GFP expression level schematic and images representing weak, moderate, and strong GFP expression, b) Weak fat body driver lines. c) Moderate fat body driver lines. d) Strong fat body driver lines. Endogenous GFP (green) and nuclei/DAPI (blue). Scale bars, 50 μm.

### Fat body driver lines induce expression in the gut

For the 27 Gal*4* lines that drove expression in the adult fat body, we next examined whether expression was also induced in the adult gut. The adult *D. melanogaster* gut is comprised of regionally distinct sections, including the proventriculus (cardia), midgut, and hindgut (Capo et al., 2013). For each *Gal4* line, we assessed GFP reporter expression in each region. Of the 27 fat body drivers, 22 had observable GFP expression in the gut (Table 2) (Fig. 3, Fig. S5, S6). We categorized these lines based on their range of expression throughout the gut. Lines such as *Met-Gal4* and *Kr-Gal4* showed expression ubiquitously or nearly ubiquitously, in all or most cell types across all three regions examined (Fig. 3). In other lines, expression was restricted to a particular region of the gut. For example, *T155-Gal4* only induced expression in the midgut while *OK376-Gal4* induced expression in the proventriculus and hindgut (Fig. 3). Finally, several lines drove expression in all three main gut regions, but not in all cell types such as *how-Gal4* or *ppl-Gal4* (Fig. 3). Notably, several lines exhibited sexually dimorphic expression. Specifically, *c564-Gal4* drove broader expression in the male midgut compared to females. For *Lk6-Gal4*, females showed expression in the midgut and malpighian tubules, while males showed expression in the proventriculus and the midgut. Taken together, our data indicates that the adult fat body and gut share many overlapping driver lines.

**Table 2.**
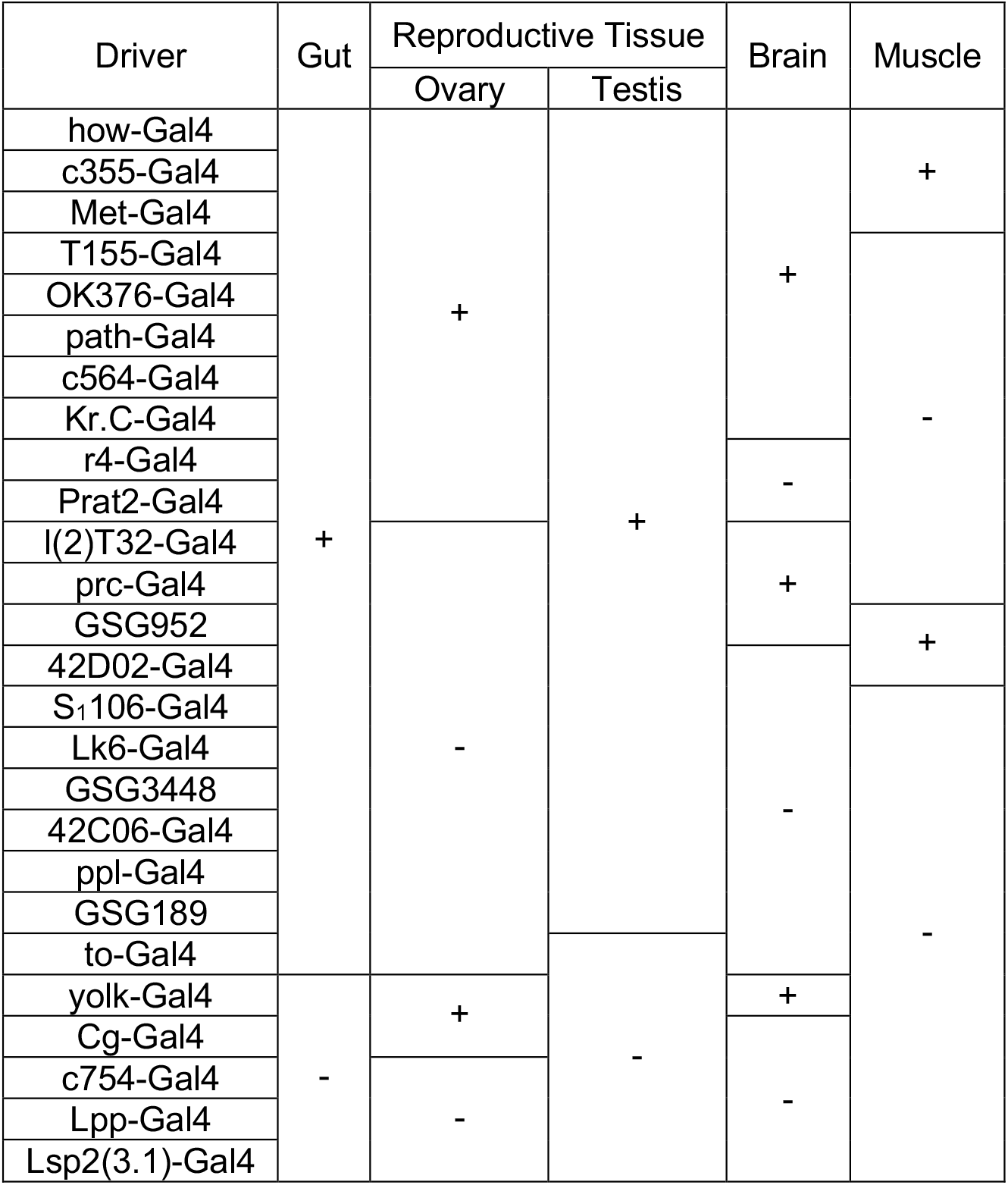
Non-fat body driver expression in adults.

**Fig. 3.**
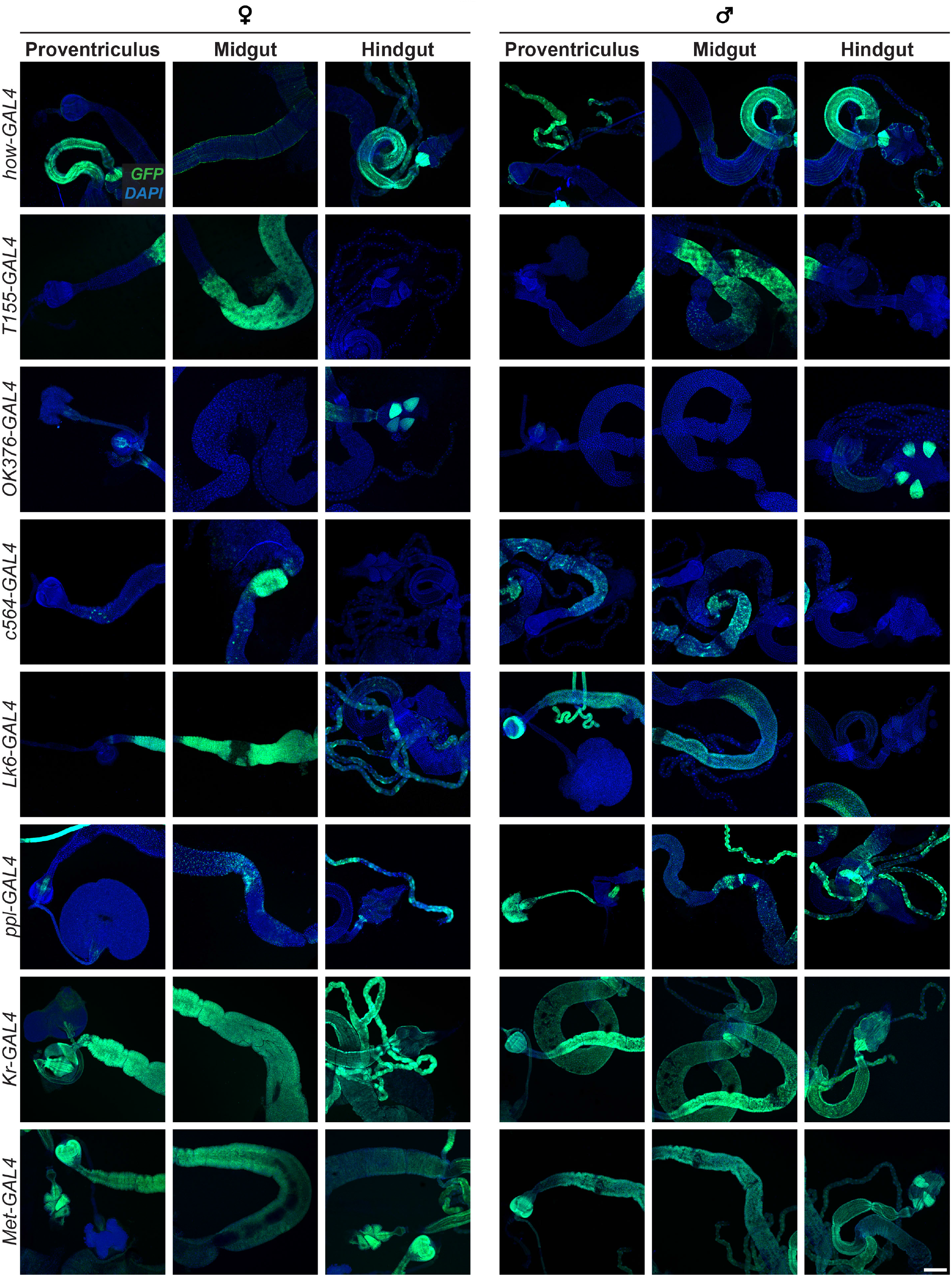
Fat body driver lines with different expression patterns within the gut. Female and male representative images of the gut. For each line, images of the proventriculus, midgut, and hindgut are shown. Endogenous GFP (green) and nuclei/DAPI (blue). Scale bar, 200 μm.

### Different fat body driver lines induce expression in the reproductive tissues

The *D. melanogaster* ovary consists of somatic and germ cells of varying types that can be targeted for genetic manipulation (Song et al., 2002). We next assessed if any fat body drivers also drove expression within the ovary. Three out of the 27 lines screened drove expression within the germaria, the most anterior region of the ovariole, or individual oocyte-producing unit of the ovary (Fig. 4a). *how-Gal4* drove expression in terminal filament and stalk cells while *yolk-Gal4* drove expression in follicle cells that encapsulate germline cysts (Fig. 4a). *Met-Gal4*, which drove broad expression throughout the ovariole, including the germarium (Fig. 4a, b). Including the three described above, 12 of the 27 lines drove expression within the ovariole, with most (11 of 12) inducing expression in ovarian follicle cells. These driver lines induced different expression patterns in ovarian follicle cell populations. The general trend for these ovarian drivers was expression in follicle cells, similar to that observed with *c355-*gal4 or T155*-Gal4* (Fig. 4b). In contrast, *yolk-Gal4* induced expression in only follicle cells that had migrated over the oocyte (Fig. 4b). Additionally, *OK376-Gal4, Kr-Gal4*, and *path-Gal4* drove expression within stretch follicle cells associated with nurse cells (Fig. 4b). *c564-Gal4* and *Prat2-Gal4* both drove expression in follicle cells bordering the nurse cells and oocytes in late-stage egg chambers. Expression in *c564-Gal4* appeared to be specific to centripetally migrating follicle cells (Fig. 4b). Finally, within the ovary, *r4-Gal4* drove expression in late dorsal-anterior follicle cells (Fig. 4b).

**Fig. 4.**
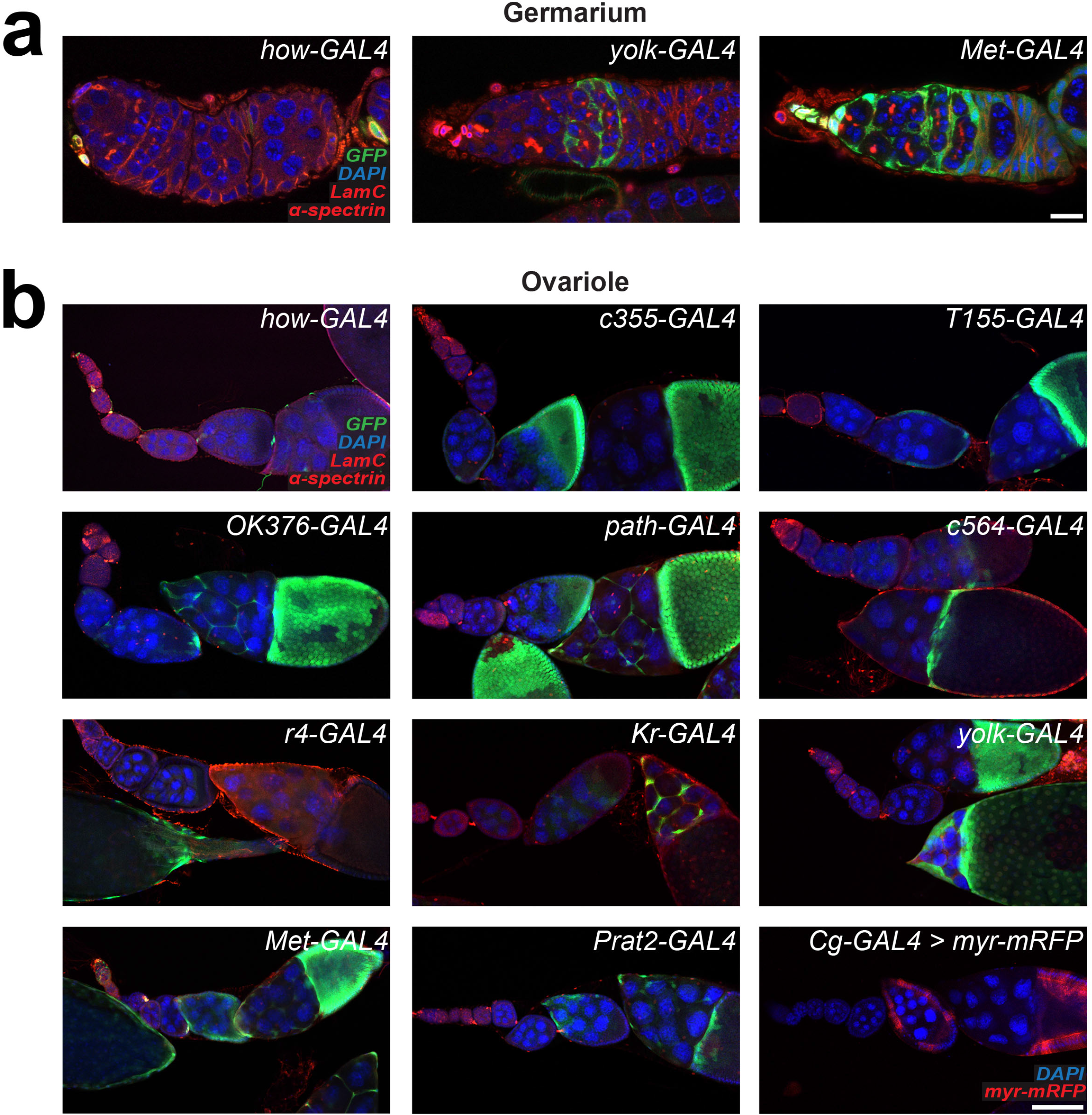
Expression patterns of fat body driver lines within the ovary. a) *Gal4* line expression patterns in the germarium. b) *Gal4* lines expression patterns in the ovariole. Labeled with LamC and alpha spectrin (red, nuclear lamina and cell membranes), endogenous GFP (green), and nuclei/DAPI (blue). Scale bar, 100 μm (germarium); 10 µm (ovariole).

In the *D. melanogaster* male reproductive system, the testis, seminal vesicle, accessory glands, and the ejaculatory duct and bulb work together to produce, maintain, and release sperm (Markow et al., 2007). To determine whether fat body driver lines induced expression in the male reproductive system, we examined each male reproductive component for GFP expression. 21 out of 27 drove expression in at least one reproductive component (Table 2) (Fig. 5, Fig. S7, S8). Lines including *OK376-Gal4, Met-Gal4*, and *prc-Gal4* had observable expression within the testis itself (Fig. 5). Other lines show broad expression throughout the reproductive system. These lines included *Met-Gal4*, driving expression in every component, and *Lk6-Gal4*, driving expression in every component except the testis (Fig. 5). Together, our analysis shows that both the female and male reproductive systems share several driver lines with the adult fat body.

**Fig. 5.**
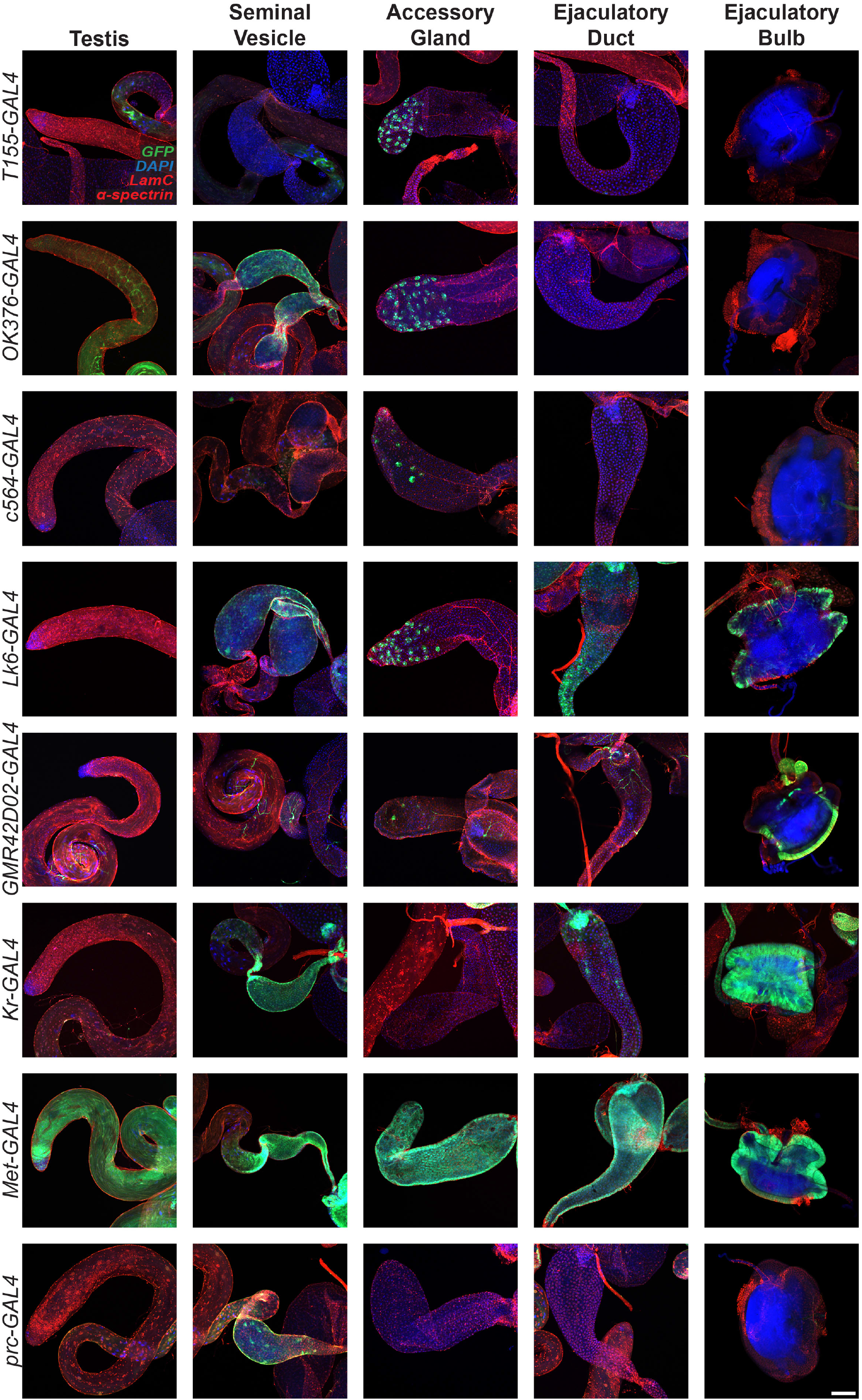
Expression patterns of fat body driver lines within the male reproductive system. *Gal4* line expression patterns in the male reproductive components. A representative image is shown for the testis, seminal vesicle, accessory gland, ejaculatory duct, and bulb. Labeled with LamC and alpha spectrin (red, nuclear lamina and cell membranes), endogenous GFP (green), nuclei/DAPI (blue). Scale bar, 100 μm.

### Different fat body driver lines induce expression in the brain and muscle

The *D. melanogaster* brain comprises multiple regions that control distinct organismal functions, including the optic lobes for visual processing, the antennal lobes for olfactory processing, and the mushroom bodies for learning and memory (Ito et al., 2014; Robinson et al., 2023; Shinomiya et al., 2022; Williams et al., 2022; Davis, 2023). We next asked whether fat body drivers induce reporter expression in the brain. 13 of 27 fat body driver lines induced expression in the brain in specific cellular subtypes, as evidenced by their location within different regions. For example, *OK376-Gal4* drove expression in the suboesophageal ganglia, *c564-Gal4* drove expression in the antennal lobes, and *yolk-Gal4* drove expression in the optic lobes (Fig. 6). Other lines, such as *Kr-Gal4* and *Met-Gal4*, exhibited broader expression throughout the brain (Fig. 6). While most of the fat body driver lines did not have any observable expression in the brain, our analysis indicated that expression induced in the brain varies for each driver line.

**Fig. 6.**
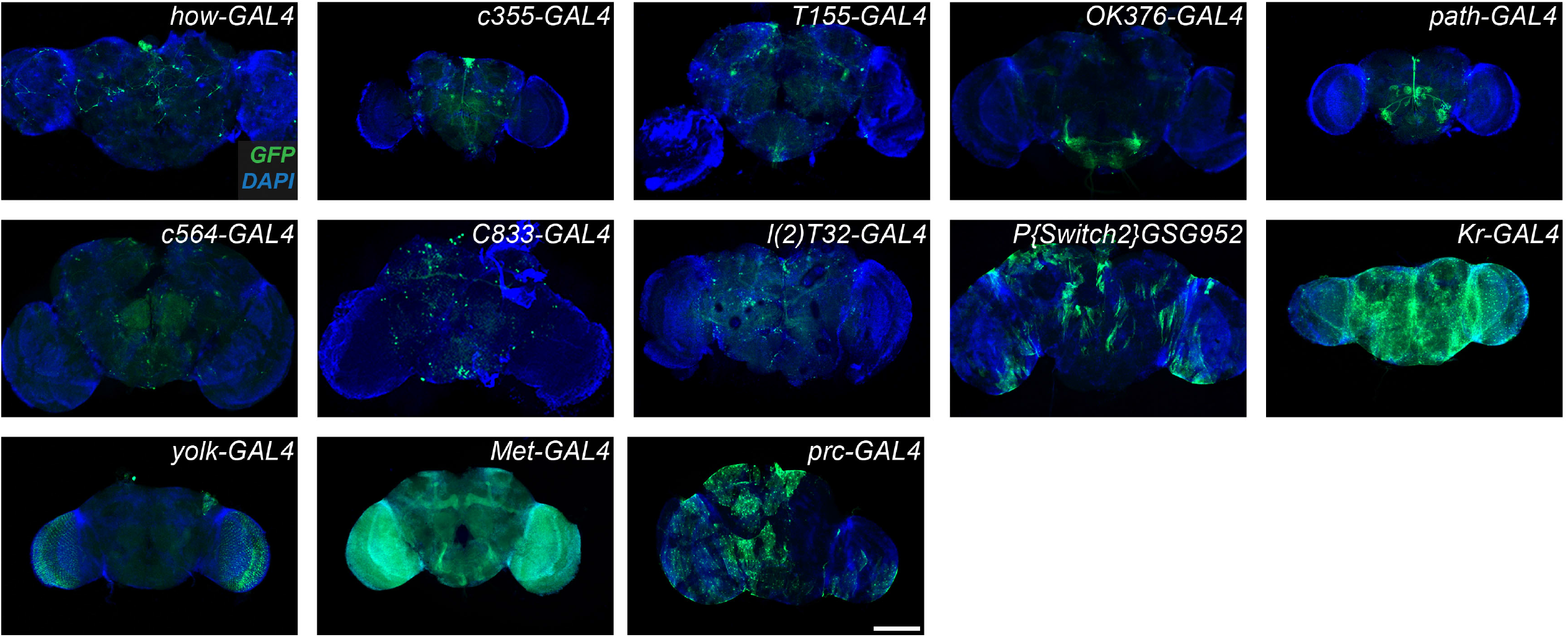
Expression patterns induced by fat body driver lines in the brain. Expression patterns of *Gal4* lines in the brain. Scale bar, 200 µm.

*D. melanogaster* adult muscle supports flight and movement (Laurichesse & Soler, 2020). We next asked if GFP expression was induced by fat body driver lines in the thoracic and abdominal muscles. Five of 27 fat body driver lines induced expression in muscles, with variability in expression patterns. *c355-Gal4, Met-Gal4, GMR42D02-Gal4*, and *P{Switch2}GSG952* drove expression in the muscle within the thorax, while *how-Gal4* drove expression in abdominal muscle (Fig. 7). Our analysis indicates that the adult fat body and muscle share fewer overlapping driver lines compared to the other tissues screened.

**Fig. 7.**
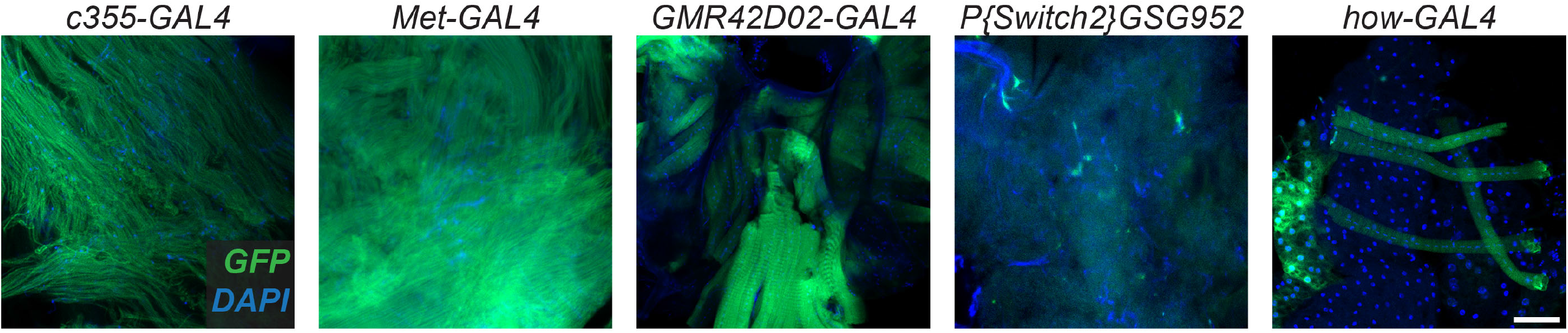
Expression patterns induced by fat body driver lines in the muscle. Expression patterns of *Gal4* lines in the muscle. Scale bar, 50 µm.

## Discussion

Transgenic *D. melanogaster Gal4* lines allow tissue- and cell type-specific control of gene expression; however, under-characterized expression patterns can lead to confounding results and data misinterpretation. In this study, we thoroughly characterized the expression patterns of 31 fat body Gal4. We observe variation in fat body Gal4 driver expression between larvae and adults (Fig. 1; Table 1), as well as between adult male and female tissues (Fig. 1 and 3; Table 1). Additionally, we find that fat body Gal4 lines vary in the expression level to induced (Fig. 2; Table 1). Importantly, we show that many of these Gal4 lines drive expression in previously undescribed secondary sites.

The strength of a Gal4 line is an important parameter when designing an experiment. In this report, we demonstrated that reported fat body drivers can induce transgene expression at different levels in the fat body (Table 3). We characterized lines, including *c754-Gal4* and *ppl-Gal4*, that exhibited strong reporter expression (Fig. 2d). This strong expression was hallmarked by bright fluorescence throughout the cytoplasm and fluorescent nuclear enrichment. Lines including *how-Gal4* and *C833-Gal4* exhibited weak reporter expression (Fig. 2b), which was characterized by dim fluorescent and sporadic expression, where GFP fluorescence was visible in some adipocytes but not in others. While we assessed variations in reporter expression, another important consideration is the amount of Gal4 protein itself. Excessive Gal4 protein has been shown to have unintended detrimental effects, including apoptosis in the imaginal eye discs and adult ventral lateral neurons (Kramer and Staveley, 2003; Rezával et al., 2007) or increased susceptibility to infection in the fat body (Keith et al., 2026). With this report, the fat body Gal4-driven reporter expression levels described can be used as a resource for selecting lines that drive weak, moderate, or strong transgene expression.

Organismal physiology impacts gene expression (Newell et al., 2020; Rivera et al., 2020; Tan et al., 2025), which in turn may lead to variations in expression patterns across Gal4 drivers. For example, neuronal and glial Gal4 drivers show reduced expression in the aging *Drosophila* brain (Delandre et al., 2025), and adult female flies fed a protein-poor diet abolish Lsp2-Gal4-driven reporter expression in the fat body (Armstrong et al., 2014). In this study, we further demonstrate differences in fat body driver expression in response to life stage, larval versus adult, and sex (Fig. 1a and b). These expression differences due to organismal physiology can be utilized to limit gene expression to a particular sex or life stage. Potential lines include yolk*-Gal4* targeting the adult female fat body, 42*D02-Gal4* and *to*-*Gal4* targeting the adult fat body, and *c729-Gal4, T110-Gal4, en2*.*4-Gal4*, and *P{Switch2}GSG10751* targeting the larval fat body. Taken together, testing whether experimental conditions affect Gal*4* expression is an important factor of consideration when incorporating a *Gal*4 line into an experiment.

Our screen identified *Lsp2(3*.*1)-Gal4* and *Lpp-Gal4* as lines that are exclusively expressed in the adult fat body (Table 2). Both *Lsp2(3*.*1)-Gal4* and *Lpp-Gal4* use promoters of fat body-derived proteins, which allows for their specificity. Rich in aromatic amino acids, Larval serum protein 2 (Lsp2) functions as a storage protein (Beneš et al., 1990), while the apoB-family lipoprotein Lipophorin (Lpp) functions as the major hemolymph lipid carrier, facilitating lipid transfer from the gut to the fat body (Palm et al., 2012). Occasionally used with the Gal4/UAS system is the Gal80 construct, which allows for Gal4 inhibition (McGuire et al., 2004). The use of the Gal80 system combined with the Gal4/UAS system can enable the generation of additional fat body-exclusive drivers.

This report presents an expression atlas of fat body driver lines that can serve as a resource to support future *D. melanogaster* fat body studies. Future areas for expansion include quantitatively scoring expression levels and examining how other physiological factors, such as age and diet, affect *Gal4* expression. Knowledge obtained from the *D. melanogaster* fat body can provide the scientific and health communities with greater insight into adipose tissue function under normal and disease conditions. Furthermore, research using *D. melanogaster* can help shed light on the intercellular and molecular mechanisms underlying adipose tissue’s role as a metabolic and endocrine organ.

## Data availability statement

The authors affirm that all data necessary for confirming the conclusions of the article are present within the article, figures, and tables.

## Acknowledgments

We thank the Bloomington *Drosophila* Stock Center for fly stocks. The monoclonal antibodies developed by P.A. Fisher, D. Branton, and R. Dubreuil were obtained from the Developmental Studies Hybridoma Bank, created by the NICHD of the NIH and maintained at The University of Iowa, Department of Biology, Iowa City, IA 52242.

